# A comparison of the Mouse Song Analyzer and DeepSqueak ultrasonic vocalization analysis systems in C57BL/6J, FVB.129, and FVB mice

**DOI:** 10.1101/2021.03.17.435868

**Authors:** Matthew S. Binder, Zachary P. Pranske, Joaquin N. Lugo

## Abstract

Vocal communication is an essential behavior in mammals and is relevant to human neurodevelopmental conditions. Mice produce communicative vocalizations, known as ultrasonic vocalizations (USVs), that can be recorded with various programs. The Mouse Song Analyzer is an automated analysis system, while DeepSqueak is a semi-automated system. We used data from C57BL/6J, FVB.129, and FVB mice to compare whether the DeepSqueak and Mouse Song Analyzer systems measure a similar total number, duration, and fundamental frequency of USVs. We found that the two systems detected a similar quantity of USVs for FVB.129 mice (*r*= .90, *p*< .001), but displayed lower correlations for C57BL/6J (*r*= .76, *p*< .001) and FVB mice (*r*= .60, *p*< .001). We also found that DeepSqueak detected significantly more USVs for C57BL/6J mice than the Mouse Song Analyzer. The two systems detected a similar duration of USVs for C57BL/6J (*r*= .82, *p*< .001), but lower correlations for FVB.129 (*r*= .13, *p*< .001) and FVB mice (*r*= .51, *p*< .01) were found, with DeepSqueak detecting significantly more USVs per each strain. We found lower than acceptable correlations for fundamental frequency in C57BL/6J (*r*= .54, *p*< .01), FVB.129 (*r*= .76, *p*< .001), and FVB mice (*r*= .07, *p*= .76), with the Mouse Song Analyzer detecting a significantly higher fundamental frequency for FVB.129 mice. These findings demonstrate that the strain of mouse used significantly affects the number, duration, and fundamental frequency of USVs that are detected between programs. Overall, we found that DeepSqueak is more accurate than the Mouse Song Analyzer.

## Introduction

The production of vocalizations plays a significant role in mammalian communication. Mammals ranging from mice to humans emit vocalizations to express social, emotional, reproductive, and ecologically relevant information ^1–5^. Alterations in vocalizations reveal insights into an animal’s health and characterize certain disorders. For instance, alterations in communication have been reliably shown to be a behavioral marker of Autism spectrum disorder (ASD), tuberous sclerosis complex (TSC), and epilepsy, in both human populations ^6–9^ and in preclinical models ^8–12^. Altogether, the production of vocalizations has many essential functions and is highly applicable to human health.

Murine models have been used to better understand the implications and applications of vocal communication. In mice, vocal communication occurs in the ultrasonic range, which exceeds the human auditory spectrum, and is subsequently referred to as ultrasonic vocalizations (USVs) ^13^. USVs are emitted between 30-90 kHz and occur in a variety of social contexts including: mother infant relationships, male-female socio-sexual interactions, and in same-sex social encounters, comprising a robust behavior that occurs throughout the lifespan^14–16^.

There are numerous automated and semi-automated programs that have been developed to assess ultrasonic vocalizations. One such program is the Mouse Song Analyzer (v.1.3), developed by the Jarvis lab (http://jarvislab.net/research/mouse-vocal-communication/). The Mouse Song Analyzer is a widely available system used in numerous publications ^17–22^. It is fully automated and extremely time efficient, as each file takes approximately 2-3 minutes to assess. A typical project with 4 groups of 10 animals per each group would only require 80-120 hours to complete. Recently, another vocalization program has been developed called DeepSqueak. DeepSqueak is semi-automated, requiring the operator to manually accept or reject each detected vocalization ^23^. The inclusion of a human observer theoretically increases the accuracy of DeepSqueak, however, it does take more time to complete a project, as a file takes on average 10 minutes to score. Thus, a project with 4 groups of 10 animals per each group in the DeepSqueak system, would require approximately 400 hours to complete, as opposed to the Mouse Song Analyzer’s 80-120 hours. Each program appears to have advantages and disadvantages over the other, however, the two programs have never been directly compared, thus the strengths and weaknesses of the Mouse Song Analyzer relative to those of DeepSqueak are unknown.

The lack of a comparison between DeepSqueak and the Mouse Song Analyzer is significant, as not all USV programs are equally accurate. Coffey et al. (2019) compared DeepSqueak to the fully automated Mouse Ultrasonic Profile ExTraction (MUPET) recording system as well as the Ultravox recording system and found that DeepSqueak had a lower USV miss rate and fewer false positives than either automated system ^23^. Any discrepancy between USV systems could lead to results that are not consistent across programs. The current study directly compared the DeepSqueak analysis program to the Mouse Song Analyzer’s analysis program using USVs collected from C57BL/6J, FVB.129, and FVB male and female mice at different ages to assess if the quantity and spectral characteristics of USVs detected in one program are similar to the quantity and spectral characteristics of USVs detected in the other.

## Materials and methods

### Animals and housing

Data from C57BL/6J, FVB.129, and FVB mice were used in the current study to evaluate the accuracy of DeepSqueak when compared to the Mouse Song Analyzer. These particular strains were assessed to give our study broad generalizability, since C57BL/6J, FVB.129, and FVB mice comprise 52% of the most popular mouse strains purchasable from Jackson Lab ^24^(Jackson Lab). USVs were collected from C57BL/6J neonates on postnatal day (PD) 12, (sample size: males = 15, females =15), PD 8 FVB.129 mice (sample size: males = 15, females = 9) and in PD 11 mice on an FVB based background (sample size: male = 12, female = 11). Altogether, this gave us a final *n* of 77 data files. All neonates used in this study were group housed with their respective dam. Across all experiments, animals had *ad libitum* access to food and water. All procedures performed were in accordance with Baylor University’s Institutional Animal Care and Use Committee, as well as the National Institutes of Health Guidelines for the Care and Use of Laboratory Animals. All USVs were recorded during the light cycle, specifically between 8 am and 5 pm.

### Isolation-induced ultrasonic vocalizations

The vocalizations were elicited via the isolation-induced ultrasonic vocalization paradigm, which has been previously described ^11,25^. Briefly, 30 minutes prior to assessment, pups were brought down from the colony and were habituated to the testing room. After 30 minutes of habituation, the pups were removed from their parents and placed into a clean holding cage that was warmed to an ambient nesting temperature of approximately 35° C. The pups were then individually removed from the holding cage and placed into a clean testing cage within a 40 cm × 40 cm *×* 30 cm acrylic, sound-attenuated chamber. All ultrasonic vocalizations emitted were recorded with a broad-spectrum condenser microphone (CM16/CMPA, Avisoft Bioacoustics, Germany) connected to a recording interface (UltraSoundGate 116Hb, Avisoft Bioacoustics). The microphone’s range was 1-125 kHz, encompassing the full ultrasonic range of the mice. Each mouse was assessed for a 2-minute duration. Once all mice were tested, they were returned to their dam in the home cage.

### MATLAB Mouse Song Analyzer analysis

The MATLAB Mouse Song Analyzer analysis procedure has been previously described ^17, 18^. Specifically, vocalization .wav files were converted to sonograms using MATLAB syntax that was developed by the Jarvis lab (http://jarvislab.net/research/mouse-vocalcommunication/). The sonograms were then processed using the graphical user interface with a density inter-syllable interval (ISI) of 0.25, a minimum frequency of 15,000 Hz, a max frequency of 125,000 Hz, a sampling frequency of 250 kHz, and a threshold of 0.3. These parameters align with previous studies and are comparable to DeepSqueak’s detection parameters ^17, 18^.

### DeepSqueak analysis

The DeepSqueak analysis system has also been previously described ^23^. Briefly, DeepSqueak was downloaded from Github via the following link (https://github.com/DrCoffey/DeepSqueak) and accessed with MATLAB 2018a. DeepSqueak has one general purpose network, one network for mouse (adult) USVs, one network for short rat USVs, and one network for long 22 kHz rat USVs. The short rat call network was used in the present study, as it had the highest correlation in total USVs detected for our mouse neonatal USV files. Once the network was established, the .wav files were imported into DeepSqueak and the following parameters for the files were set: a total analysis length of 0, an analysis chunk length of 6, a frame overlap of .0001 seconds, a frequency low cut off of 30 kHz, a frequency high cut off of 120 kHz, and a score threshold of 0. The detection setting was set to “high recall” to ensure detection of as many USVs as possible. The files were then manually processed, with the tonality threshold being adjusted to optimize the signal-to-noise ratio for each file, and automatic detection boxes were redrawn as needed to capture the entire duration and frequency range of each vocalization.

### Statistical analysis

All data was analyzed using IBM SPSS Statistics 21.0 (IBM, USA) or GraphPad Prism 7 software (La Jolla, CA). The dependent variables were the total quantity, average duration, and average fundamental frequency of USVs emitted, as each of these has been shown to be of particular relevance to neurodevelopmental conditions ^8,9,11,26–28^. Correlations for each strain were performed to determine the accuracy of the Mouse Song Analyzer’s automated processing to the DeepSqueak program for each dependent variable. Additionally, to examine if DeepSqueak and the Mouse Song Analyzer detected significant differences from one another per each dependent variable, independent *t*-tests or nonparametric Mann-Whitney *U* tests (when homogeneity of variance was violated) per each strain were run. A partial correlation analysis was then conducted to control for the following covariates: sex (male, female), strain (C57BL/6J, FVB.129, FVB), and age (PD 8,11,12) of the mice. Furthermore, a correlation that did not control for any of the covariates (a zero-order correlation) was also calculated to determine the influence the covariates had on the relationship in USV detection between DeepSqueak and the Mouse Song Analyzer. A value of *p*< .05 was considered significant for each statistical test and the figures depict the mean ± standard error of the mean (SEM). The effect sizes for the correlations are as follows: small effect r = 0.1, medium effect *r*= 0.3, large effect *r* = 0.5.

## Results

### Association and number of USVs detected between DeepSqueak and the Mouse Song Analyzer

For the C57/BL6J strain, there was a large correlation between the total number of USVs detected by DeepSqueak and the total USVs detected by the Mouse Song Analyzer (*r* (66) = .76, *p* < .001) (Figure 1A). A Mann-Whitney *U* test determined that DeepSqueak detected significantly more vocalizations than the Mouse Song Analyzer for C57BL/6J mice (*U* = 241.5, *p* < .001) (Figure 1B). For the FVB.129 strain, there was a large correlation in the USVs detected between the two programs (*r* (46) = .90, *p* < .001) (Figure 1C). There was also no significant difference in the quantity of vocalizations detected between DeepSqueak and the Mouse Song Analyzer for FVB.129 mice (*t* (46) = .51, *p* = .61) (Figure 1D). In the FVB strain, there was a large correlation between DeepSqueak and the Mouse Song Analyzer for the total USVs that were detected (*r* (38) = .60, *p* < .001) (Figure 1E). No significant differences in the quantity of USVs were detected between DeepSqueak and the Mouse Song Analyzer for FVB mice (*t* (44) = .38, *p* = .71) (Figure 1F).

**Figure 1.**
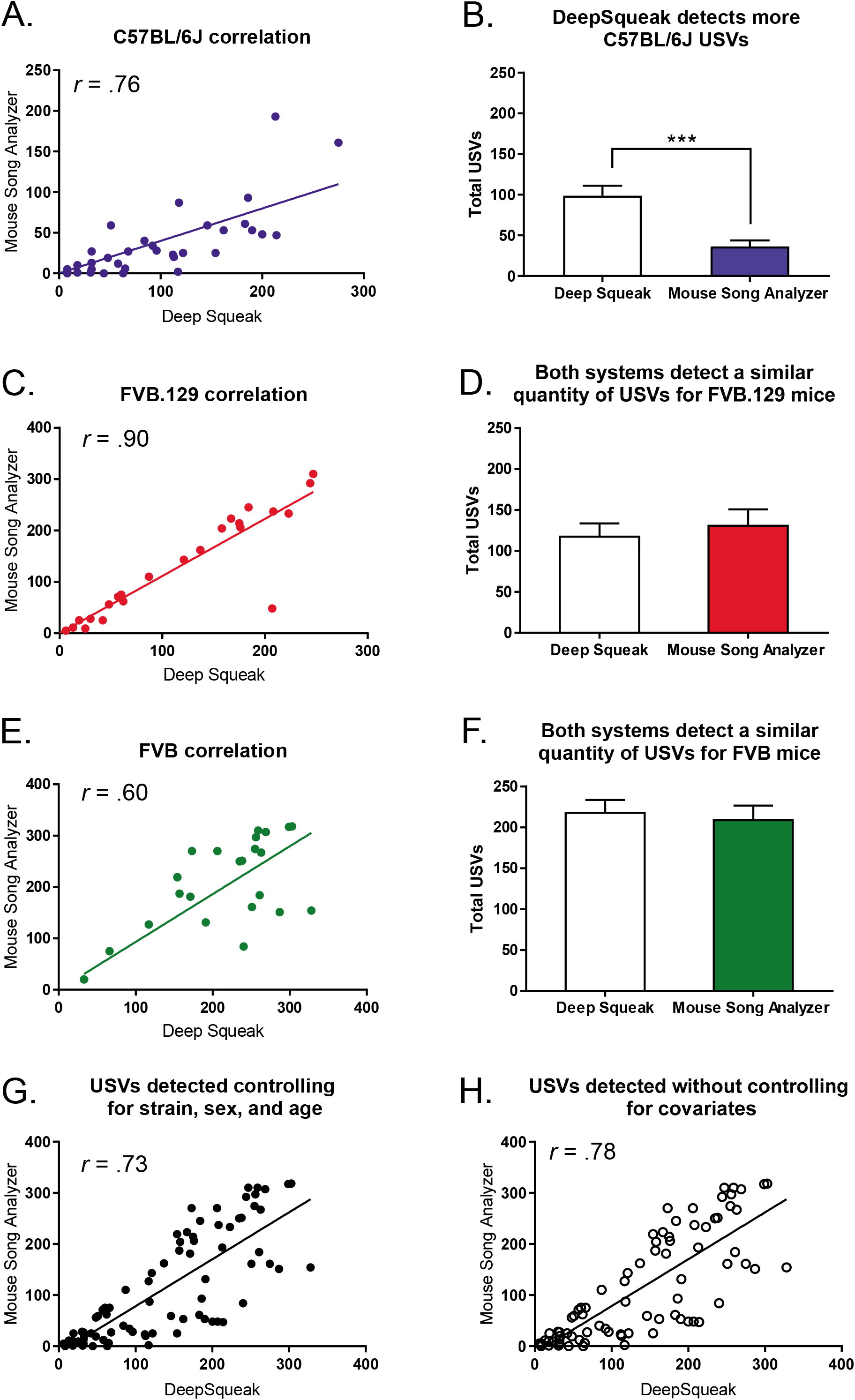
There were high correlations in USV detection between DeepSqueak and the Mouse Song Analyzer for each strain but DeepSqueak detected significantly more USVs than the Mouse Song Analyzer for C57BL/6J mice. (A). There was a large correlation between systems for C57BL/6J mice. (B). DeepSqueak detected significantly more USVs than the Mouse Song Analyzer for C57BL/6J mice. (C). There was a large correlation between systems for FVB.129 mice. (D). There was no significant difference in the quantity of calls detected between DeepSqueak and the Mouse Song Analyzer for FVB.129 mice. (E). There was a large correlation between systems for the FVB mice. (F) There was no significant difference in the quantity of calls detected between DeepSqueak and the Mouse Song Analyzer for FVB mice. (G). Upon controlling for the variables of strain, sex, and age, a large correlation was found. (H). The strain of mouse assessed was found to substantially influence the relationship between USVs detected in DeepSqueak and those detected in the Mouse Song Analyzer, whereas the sex, and age variables do not substantially influence this relationship. The data points represent the mean and the error bars represent the standard error of the mean. * = *p* < .05, ** = *p* < .01, *** = *p* < .001.

### Correlation controlling for variables of assessment for the quantity of USVs detected

A partial correlation was performed to determine the strength of the relationship between the number of USVs detected by the two systems whilst controlling for strain (C57BL/6J, FVB.129, FVB), sex (male, female), and age of the animal (PD 8,11,12). There was a large positive correlation between the number of USVs detected across the two systems, which was statistically significant (*r* (72) = 0.73, *p* < 0.001) (Figure 1G). However, the zero-order correlation was notably different (*r* (72) = 0.78, *p* < 0.001), with strain accounting for .44 of the variance (Figure 1H). Therefore, while the sex and age covariates have little influence on USV detection between systems (.15 and .3 respectively), the strain of the mouse does influence the relationship between USVs detected in DeepSqueak and those detected in the Mouse Song Analyzer.

### Average duration of USVs detected between DeepSqueak and the Mouse Song Analyzer

When comparing the average duration of the USVs detected in DeepSqueak to the Mouse Song Analyzer, we found a large correlation between systems for the C57BL/6J strain (*r* (58) = .82, *p* < .001) (Figure 2A). However, DeepSqueak detected a significantly higher average USV duration than the Mouse Song Analyzer (*U* = 48, *p* < .001) (Figure 2B). For the FVB.129 strain, there was a small correlation for the duration of the USVs detected between the two programs (*r* (46) = .13, *p* < .001) (Figure 2C). Additionally, DeepSqueak detected a significantly higher average duration of vocalizations compared to the Mouse Song Analyzer (*t* (46) = 7.96, *p* < .001) (Figure 2D). In the FVB strain, there was a large correlation between DeepSqueak and the Mouse Song Analyzer for the average duration of the USVs (*r* (38) = .51, *p* = .01) (Figure 2E). DeepSqueak detected a higher average duration of USV compared to the Mouse Song Analyzer for FVB mice (*t* (44) = 11.06, *p* < .001) (Figure 2F).

**Figure 2.**
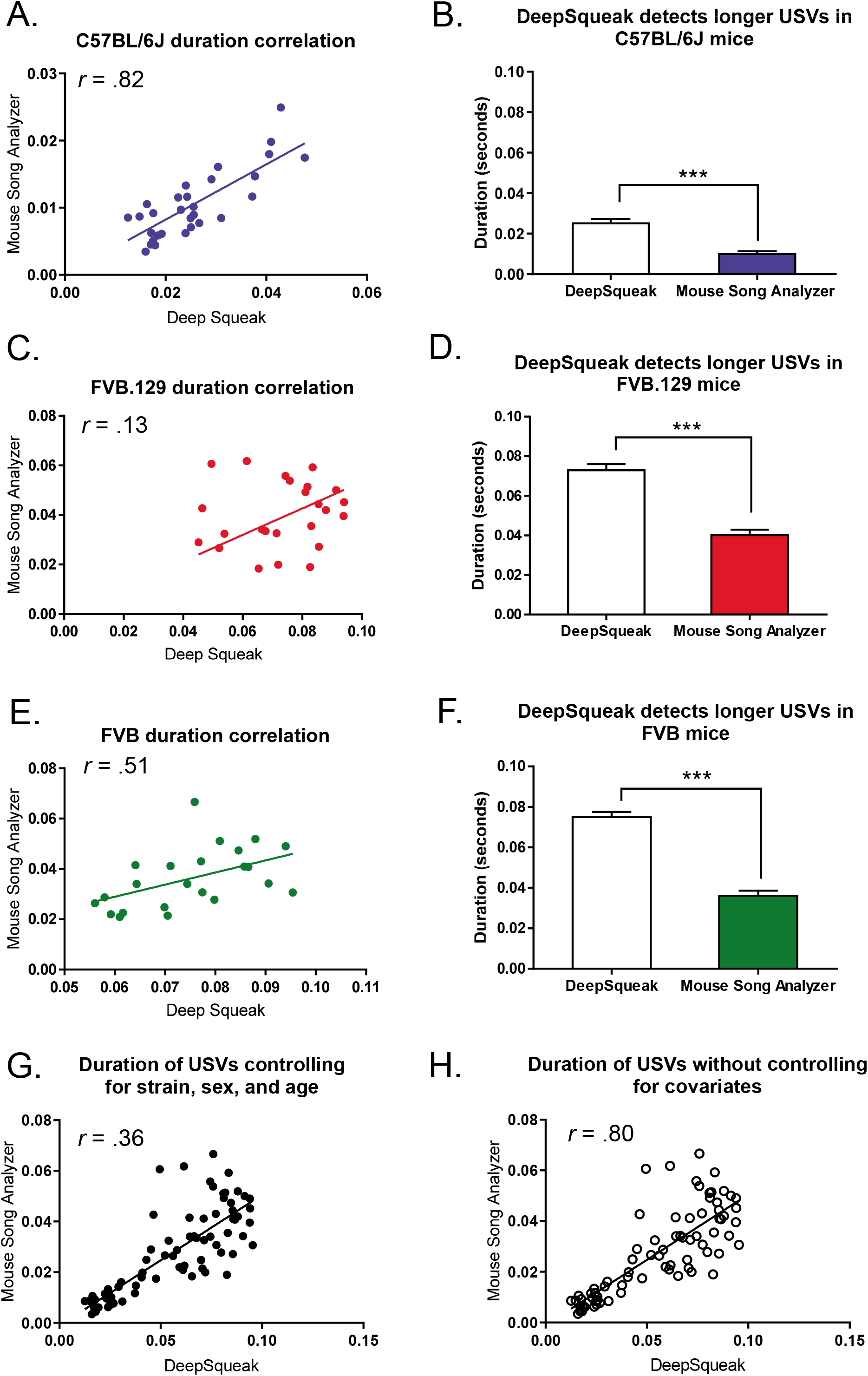
There were low to high correlations for the duration of USVs detected in DeepSqueak and those detected in the Mouse Song Analyzer for the C57BL/6J, FVB.129, and FVB strains, as well as significant differences in duration between each system. (A). There was a large correlation between systems for the duration of C57BL/6J mice. (B). DeepSqueak detected significantly longer USVs than the Mouse Song Analyzer for C57BL/6J mice. (C). There was a small correlation between systems for FVB.129 mice. (D). DeepSqueak detected significantly longer USVs than the Mouse Song Analyzer for FVB.129 mice. (E). There was a large correlation between systems for FVB mice. (F) DeepSqueak detected significantly longer USVs than the Mouse Song Analyzer for FVB mice. (G). Upon controlling for the variables of strain, sex, and age, a medium correlation was found. (H). The age of the mouse assessed was found to substantially influence the relationship between the duration of USVs detected in DeepSqueak and those detected in the Mouse Song Analyzer, whereas the sex, and strain variables did not substantially influence this relationship. The data points represent the mean and the error bars represent the standard error of the mean. * = *p* < .05, ** = *p* < .01, *** = *p* < .001.

### Correlation controlling for variables of assessment for the duration of the USVs detected

A partial correlation was performed to determine the strength of the relationship between the average duration of the USVs detected by the two systems whilst controlling for strain (C57BL/6J, FVB.129, FVB), sex (male, female), and age of the animal (PD 8, 11, 12). There was a medium positive correlation between the average duration of USVs detected across the two systems, which was statistically significant (*r* (75) = 0.36, *p* = 0.002) (Figure 2G). However, the zero-order correlation was significantly larger (*r* (75) = 0.80, *p* < 0.001), with age accounting for .79 of the variance (Figure 2H). Therefore, while the strain and sex covariates have little influence on the average duration of the USVs detected between systems (.1 and .12 respectively), the age of the mouse does influence the relationship between DeepSqueak the Mouse Song Analyzer for the duration of the vocalizations.

### Average fundamental frequency of USVs detected between DeepSqueak and the Mouse Song Analyzer

When assessing the average fundamental frequency of the USVs detected in DeepSqueak with the Mouse Song Analyzer, we found a large correlation between systems for the C57BL/6J strain (*r* (58) = .54, *p* < .01) (Figure 3A). No significant differences in the average fundamental frequency detected were found between DeepSqueak and the Mouse Song Analyzer (*t* (58) = .94, *p* = .35) (Figure 3B). For the FVB.129 strain, there was a large correlation for the fundamental frequency of the USVs detected between the two programs (*r* (46) = .76, *p* < .001) (Figure 3C). However, the Mouse Song Analyzer detected a higher fundamental frequency (*t* (46) = 2.31, *p* = .03) (Figure 3D). In the FVB model, there was no significant correlation between DeepSqueak and the Mouse Song Analyzer for the average fundamental frequency of the USVs (*r* (38) = .07, *p* = .76) (Figure 3E). Additionally, no significant difference was found in the average fundamental frequency of USVs detected between DeepSqueak and the Mouse Song Analyzer for FVB mice (*t* (44) = 1.35, *p* = .18) (Figure 3F).

**Figure 3.**
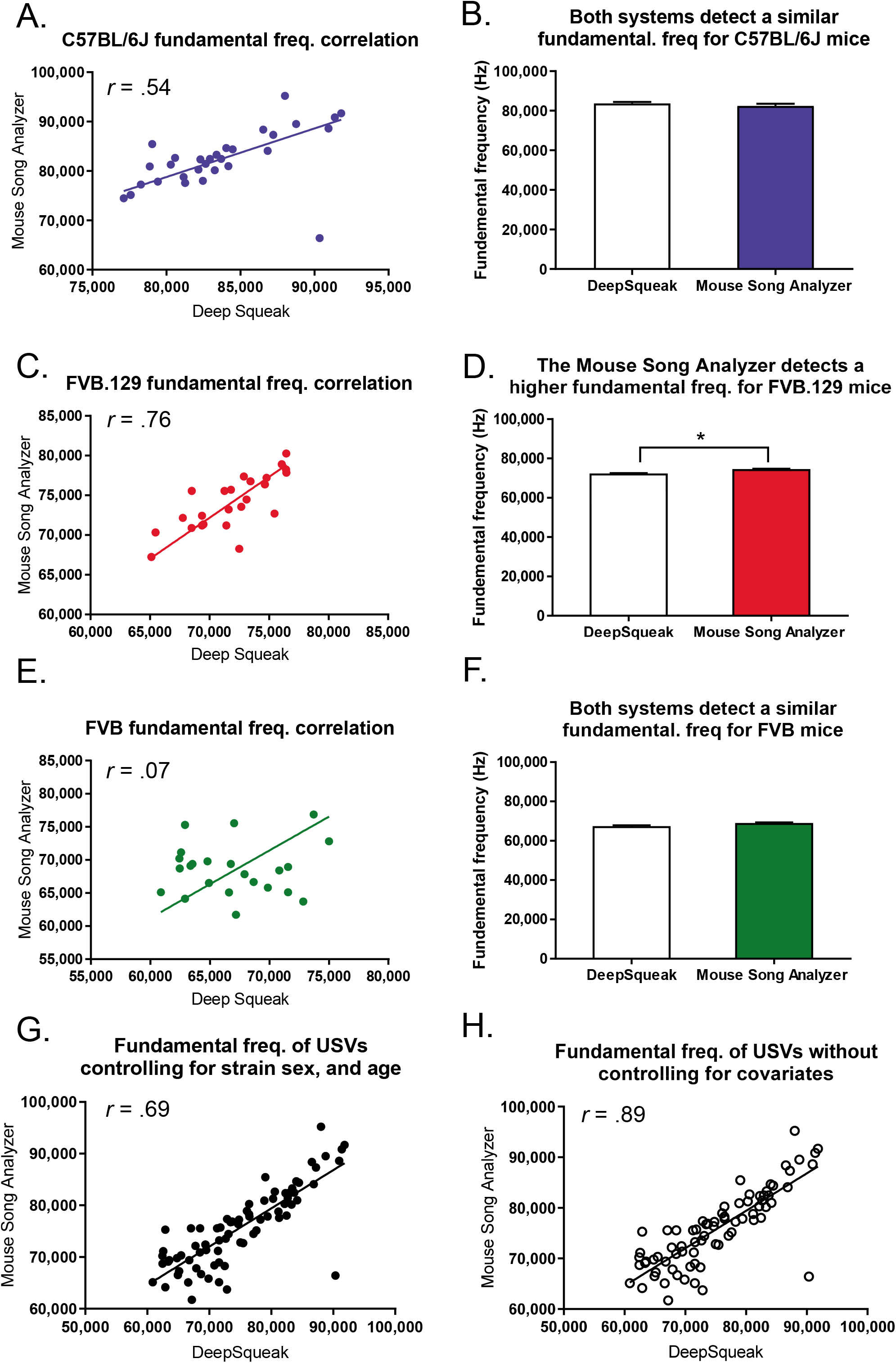
There were low to high correlations for the fundamental frequency of USVs detected in DeepSqueak and those detected in the Mouse Song Analyzer for the C57BL/6J, FVB.129, and FVB strains, and a significant difference in the fundamental frequency between each system for FVB.129 mice. (A). There was a large correlation between systems for the fundamental frequency of C57BL/6J mice. (B). There was no significant difference in the fundamental frequency of USVs detected between DeepSqueak and the Mouse Song Analyzer for C57BL/6J mice. (C). There was a large correlation between systems for FVB.129 mice. (D). DeepSqueak detected a significantly lower fundamental frequency of USV than the Mouse Song Analyzer for FVB.129 mice. (E). There was no significant correlation between systems for the FVB mice. (F) There was no significant difference for the fundamental frequency of USVs detected between DeepSqueak and the Mouse Song Analyzer for FVB mice. (G). Upon controlling for the variables of strain, sex, and age, a large correlation was found. (H). It was found that the age of the mice influenced the relationship between DeepSqueak and the Mouse Song Analyzer for the fundamental frequency parameter the most, followed by the strain variable, with the sex variable not substantially influencing this relationship. The data points represent the mean and the error bars represent the standard error of the mean. * = *p* < .05, ** = *p* < .01, *** = *p* < .001.

### Correlation controlling for variables of assessment for the fundamental frequency of the USVs detected

A partial correlation was performed to determine the strength of the relationship between the average fundamental frequency of the USVs detected by the two systems whilst controlling for strain (C57BL/6J, FVB.129, FVB), sex (male, female), and age of the animal (PD 8,11,12). There was a large positive correlation between the average duration of USVs detected across the two systems, which was statistically significant (*r* (72) = 0.69, *p* < 0.001) (Figure 3G). However, the zero-order correlation was significantly larger (*r* (72) = 0.89, *p* < 0.001), with age accounting for .73 of the variance, strain accounting for .28 of the variance and sex accounting for .05 of the variance (Figure 3H). Therefore, while the sex covariate had little influence on the average fundamental frequency of the USVs detected between systems, the age, and to a lesser extent the strain, variables did influence the relationship between DeepSqueak and the Mouse Song Analyzer for the fundamental frequency of the vocalizations.

## Discussion

The present study compared the DeepSqueak analysis system to the MATLAB Mouse Song Analyzer analysis system to assess the reproducibility of findings between USV programs. The C57BL/6J, FVB.129, and FVB strains were chosen since they comprise 52% of the most popular mouse strains purchasable from Jackson labs (Jackson Lab) ^24 24 24^. Our study also included male and female mice across a range of ages. For the total quantity of USVs detected, large correlations were found between DeepSqueak and the Mouse Song Analyzer for each strain, however, strain specific differences between systems were observed. When assessing the average duration and fundamental frequency of the USVs, there were lower correlations present and numerous significant differences were found between systems.

The standard for reliability for two systems under identical detection parameters is a correlation between .9-1 ^29^. While there were large correlations between DeepSqueak and the Mouse Song Analyzer for the number of USVs detected across each strain, the correlations were lower than anticipated for C57BL/6J (.76) and FVB mice (.60). If the data between two systems is less correlated, then results may not be consistent across systems. Our findings support this conclusion, as DeepSqueak detected significantly more USVs than the Mouse Song Analyzer for the C57BL/6J strain. Furthermore, strain was shown to account for significant variance between the two systems for the total quantity of USVs detected. Our data suggests that results attained in one system for either the C57BL/6J or FVB mouse strains may not be reproducible in the other system.

Although there was lower than expected correlations for C57BL/6J and FVB mice, this was not the case for FVB.129 mice. FVB.129 mice displayed a high correlation of .9, satisfying the standard for reliability. Furthermore, there was no statistically significant difference in the total quantity of USVs detected between DeepSqueak and the Mouse Song Analyzer, indicating that, for the 129-background strain, results were reproducible across systems.

The strain-dependent variability in USV detection between DeepSqueak and the Mouse Song Analyzer may best be attributed to the USV cut off parameters of the two programs. For C57BL/6J and FVB mice, the MATLAB algorithm that detects vocalizations may do so in a manner that occasionally counts 2 USVs that occur close together in time as one larger USV or may miss a vocalization if it is emitted in the presence of background noise. Conversely, the DeepSqueak program requires a human operator to verify its automated detection algorithm. The operator is presented with both visual and auditory cues for each call and must accept or reject each detected vocalization. Therefore, the semi-automated aspect of DeepSqueak reduces false positives and problems with USV clustering. Furthermore, a prior study demonstrated that in both low and high background noise conditions DeepSqueak was similarly accurate, indicating that it is adept at mitigating false negatives and correctly detecting USVs regardless of interference, thus DeepSqueak may be avoiding a confound that could be biasing the Mouse Song Analyzer’s detection algorithm ^23^. Interestingly, when assessing FVB.129 mice, we found high reliability between systems. This may be attributed to 129 mice innately vocalizing in a pattern that better conforms to the Mouse Song Analyzer’s automated detection algorithm than either the C57BL/6J or FVB strains, limiting any discrepancies in USV detection. For instance, a previous study by Wiaderkiewicz (2013), found that in a 7-minute test, 129 mice emitted fewer vocalization clusters than C57BL/6J mice. Any reduction in the clusters of vocalizations in a file may simplify the complexity of USV detection and better align with the Mouse Song Analyzer’s algorithm ^30^. Furthermore, Rogers et al. (2002) showed that 129 background mice exhibit a hypolocomotor profile when compared to the C57BL/6J background ^31^. Although Rogers et al. (2002) used adult mice, reduced locomotion during USV testing would decrease ambulatory background noise levels, limiting interference and simplifying USV detection, potentially increasing the accuracy of the Mouse Song Analyzer. Overall, several factors contribute to the differences in vocalization detection between DeepSqueak and the Mouse Song Analyzer, which may lead to inconsistencies between programs depending on the strain of mouse assessed.

We also investigated the reliability between systems for the average duration and fundamental frequency of the vocalizations. These two parameters were chosen because previous preclinical models of neurodevelopmental diseases, such as ASD and epilepsy, have found alterations in the duration and fundamental frequency of USVs, with clinical studies in ASD infants also reporting changes to these communicative features, indicating that they may be of particular pertinence to disease states and an animal’s health ^8,9,11,26–28^. Across all strains, we found that the correlations between DeepSqueak and the Mouse Song Analyzer for the duration and fundamental frequency of the USVs ranged from .07-.82. Therefore, none of the spectral and temporal characteristics of the USVs satisfied the standard for reliability across systems, regardless of the strain assessed. Follow-up statistical analyses found numerous significant differences between systems for the mean duration and fundamental frequency, supporting the correlation data analyses indicating that results attained from one program may not be similar in the other.

The significant variability between programs for the spectral and temporal features of the USVs may be a product of the increased complexity of these measurements compared to the relatively simple measurement of USV count. Several parameters must be taken into account when automatically calculating the average duration or fundamental frequency of each vocalization in a file. Since a single file may have over 100 USVs, any slight discrepancy in detection parameters between systems would become magnified and lead to significantly different results. Additionally, USV characteristics are intrinsically tied to USV detection, therefore the factors mentioned earlier, such as variability in background noise, USV clustering, and the occurrence of false positives, could also bias the spectral and temporal assessment of USVs. Importantly, this also means that DeepSqueak is a more accurate system for detecting USV call characteristics than the Mouse Song analyzer, due to its reliance on an operator to ensure the validity of each USV. Thus, the more accurate detection of USVs in DeepSqueak results in a more accurate assessment of USV’s spectral and temporal characteristics than the Mouse Song Analyzer. Since the duration and the fundamental frequency have been reported to be important markers of communication, any inconsistencies across USV programs in the detection of a call’s spectral and temporal parameters are significant and represent a compelling future direction for further study.

Previous studies have found that only 36-61% percent of the studies in psychology, behavioral and medical journals are able to be replicated ^32–34^. Since science is a cumulative endeavor that builds upon past work, any inability to reproduce prior findings can have a long-lasting and detrimental impact for research. In light of the reproducibility crisis, the NIH has initiated a charge to enhance rigor and reproducibility in science ^35^. While our study takes a step towards fulfilling this goal, more studies need to follow to ensure consistent results in USV research. Future studies could continue to compare DeepSqueak and the Mouse Song Analyzer across other commonly used mouse strains. Additionally, while USV detection in DeepSqueak has been shown to be more accurate and robust than the USV detection in the Ultravox and the Mouse Ultrasonic Profile ExTraction (MUPET) vocalization systems, the spectral and temporal characteristics of the USVs were not assessed across systems. Therefore, the reliability of the spectral features of vocalizations in different recording programs is unknown. The internal validation of vocalization methodology is of the utmost necessity to ensure the integrity and productivity of the field and will allow for USV results that are not contingent upon the software used in analysis.

## Conclusion

In the current study, we observed that not all data may be reproducible between DeepSqueak and the Mouse Song Analyzer, as correlations between systems for the C57BL/6J and FVB strains were lower than generally acceptable. Additionally, there was significant variation when assessing the duration and fundamental frequency of the USVs across programs regardless of strain. Overall, DeepSqueak was found to detect more USVs than the Mouse Song Analyzer however, the increased accuracy comes at the expense of time, as DeepSqueak takes 10-15 minutes to assess a file whereas the Mouse Song Analyzer takes only 2-3 minutes. Importantly, the Mouse Song Analyzer still yields a statistically large correlation for total USV count, indicating that the majority of USVs are detected. Overall, both systems have relative advantages and disadvantages that need be taken into consideration by vocal communication researchers when interpreting USV results or when analyzing USV files.

## Acknowledgements

The authors do not have any conflicts of interest to disclose. We would like to thank Samantha Hodges, David Narvaiz, Paige Womble, and Greg Sullens for their critical review of the paper.

## Author contributions

MSB, ZPP, and JNL conceived of the hypothesis and designed the study. MSB and ZPP performed experiments, analyzed, and plotted the data. MSB and JNL wrote the manuscript.

---

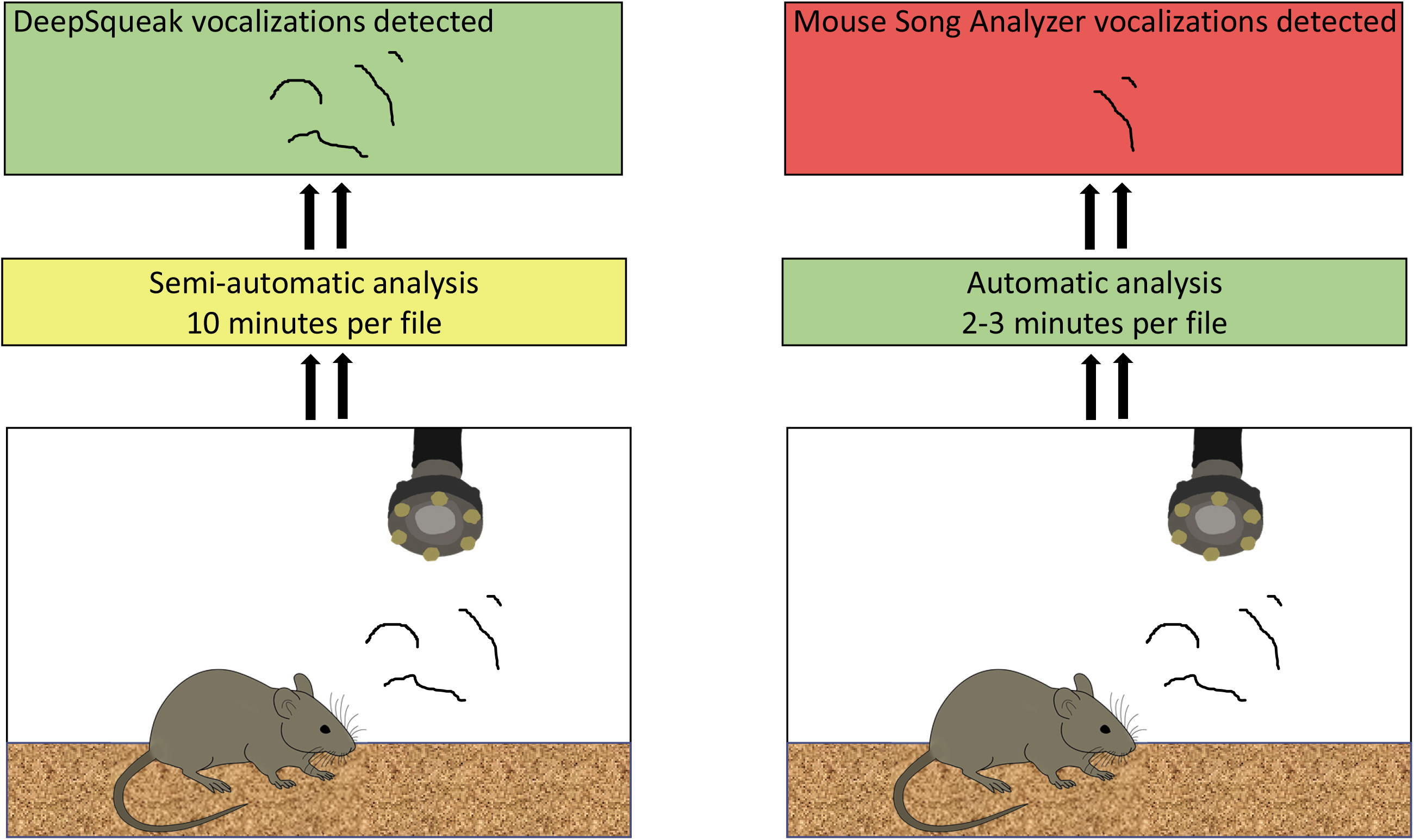

